# Distinct working-memory organisation for the colour and duration of visual objects

**DOI:** 10.64898/2026.08.31.748261

**Authors:** Anna M. van Harmelen, Freek van Ede

## Abstract

When holding information ‘in mind’, it is essential that individual objects in working memory remain separated and accessible for guiding behaviour. Space is a well-established organising principle for retaining various features of visual objects in working memory, but whether this spatial organisation is universally applied across all features of visual objects remains unclear. Here, we tested whether two inherently non-spatial features of a visual item, colour and duration, rely on the same spatial organisation of working memory. To track this spatial organisation implicitly, we measured directional biases in microsaccades while participants were cued to select one of two memorised visual items in order to report either its colour or its duration, depending on the session. Crucially, while the visual stimuli were identical between the colour-report and the duration-report sessions, accessing colour information led to a robust spatial microsaccade bias towards the item’s original location, whereas this bias was absent when accessing duration. This was corroborated by a significant difference in the spatial bias between the colour and duration sessions. We conclude that the memory organisation for colour and duration is distinct, even when memory regards the exact same visual objects. This implies that the spatial organisation of working memory for visual features is not universal, but feature-dependent.

## Introduction

A fundamental question in cognitive science is how information in working memory is organised to ensure that individual memory items can be accessed selectively for behaviour, while minimising interference with other memory items. Over the past few decades, ample studies have converged on the prevailing notion that space serves as a foundational scaffold for organising information from visual objects in working memory. When an object is encoded into working memory, its constituent features are essentially “bound” to its spatial coordinates (Awh & Vogel, 2025; Pertzov & Husain, 2014; A. Treisman & Zhang, 2006), even when location is entirely task-irrelevant and never explicitly tested. This spatial scaffolding is remarkably robust and generalises beyond geometric properties like shape (Kuo et al., 2009; A. Treisman & Zhang, 2006) or orientation (de Vries & van Ede, 2024; Draschkow et al., 2022; Poch et al., 2017; Schneegans & Bays, 2017), to purely non-spatial features such as colour (Foster et al., 2017; Kuo et al., 2009; Schneegans & Bays, 2017; Theeuwes et al., 2011; A. Treisman & Zhang, 2006; van Ede et al., 2019). Furthermore, it persists even when information is encoded sequentially and cued through serial order (de Vries et al., 2023) and after relatively long working-memory delays of at least up to five seconds (de Vries & van Ede, 2025). Collectively, these findings have fostered the prevailing view that space is a universal organising principle for any type of visual information maintained in working memory.

However, it remains unknown whether this spatial architecture is truly universal, or if certain visual properties may be exempt from such a spatial organisation. A prominent candidate feature to test the boundaries of this organisational architecture is the duration of visual items. Like colour, duration is an inherently non-spatial property of a visual stimulus. Yet, unlike other visual features, such as shape and colour, the duration of a stimulus does not exist within a single moment. Rather, it must be integrated over time and is likely processed through distinct neural networks that span beyond classical retinotopic visual cortices (Centanino et al., 2026; Coull et al., 2008; Harvey et al., 2020). While humans can reliably maintain duration information in working memory (e.g., Coull et al., 2008; Glasauer & Shi, 2021; Herbst et al., 2025; Manohar et al., 2017; Mauk & Buonomano, 2004; M. Treisman, 1963; Wassenhove et al., 2008), whether this maintenance depends on a similar spatial organisation as has been reported for other features of visual objects remains an open question.

Initial evidence that duration information in maintained in working memory without a spatial organisation comes from (Kruijne et al., 2021). They reported an absence of two classic EEG markers of spatial attention, the contralateral delay activity and lateralised alpha suppression, suggesting that location representations may indeed not be active when duration information is maintained. However, the findings from this prior study remain constrained in three main ways. First, in that study, durations were presented as the interval between brief visual flashes, rather than as a stable property of a single visual object. Second, the authors used a task with a binary comparison report, which may demand less precise memory formation than a continuous reproduction report (as is frequently used for testing other visual features like orientation or colour). Finally, without comparing duration directly to another visual feature, these earlier findings ultimately hinge on a null result. Consequently, it remains unclear whether maintaining a detailed representation of a visually stable duration operates independently of the spatial scaffolding that governs other visual features.

In the current study, we investigated whether visual durations are stored in working memory using the same spatial architecture as colours, or if duration forms an exception to the spatial-scaffolding hypothesis. To track the spatial organisation of a visual feature in working memory, we measured directional biases in microsaccades during memory recall. Specifically, participants were prompted to access one of two memorised items and report either its colour or duration, depending on the session. Critically, these biases enabled us to track the spatial organisation of memory implicitly, without ever having to ask participants about the location of memorised items (de Vries et al., 2023; de Vries & van Ede, 2024, 2025; Draschkow et al., 2022; Liu et al., 2024, 2026; for a recent overview, see also: van Ede, 2026). Using this robust implicit index of spatial memory organisation, we demonstrate a striking dissociation between the organisation of visual colour and visual duration information in working memory. Spatial microsaccade biases only emerged when recalling the colour of the cued visual memory item, and differed significantly between the colour and duration sessions. Crucially, we demonstrate this while using the exact same visual stimuli, overall task structure, and continuous reproduction reports, thereby only varying whether participants were instructed to memorise the colour or duration of the visual items for the ensuing report.

## Results

We investigated whether memorising the colour and duration of visual items rely on the same principles of spatial organisation in working memory. Healthy human volunteers performed two separate sessions differing exclusively in which visual feature of two items had to be recalled at the end of a trial: colour or duration (**Fig. 1A**). On each trial, two items were sequentially presented with randomised colours and durations. During the subsequent working-memory delay, an order cue (‘1’ or ‘2’) determined which item’s session-relevant feature had to be recalled and reproduced at the end of the trial.

**Figure 1:**
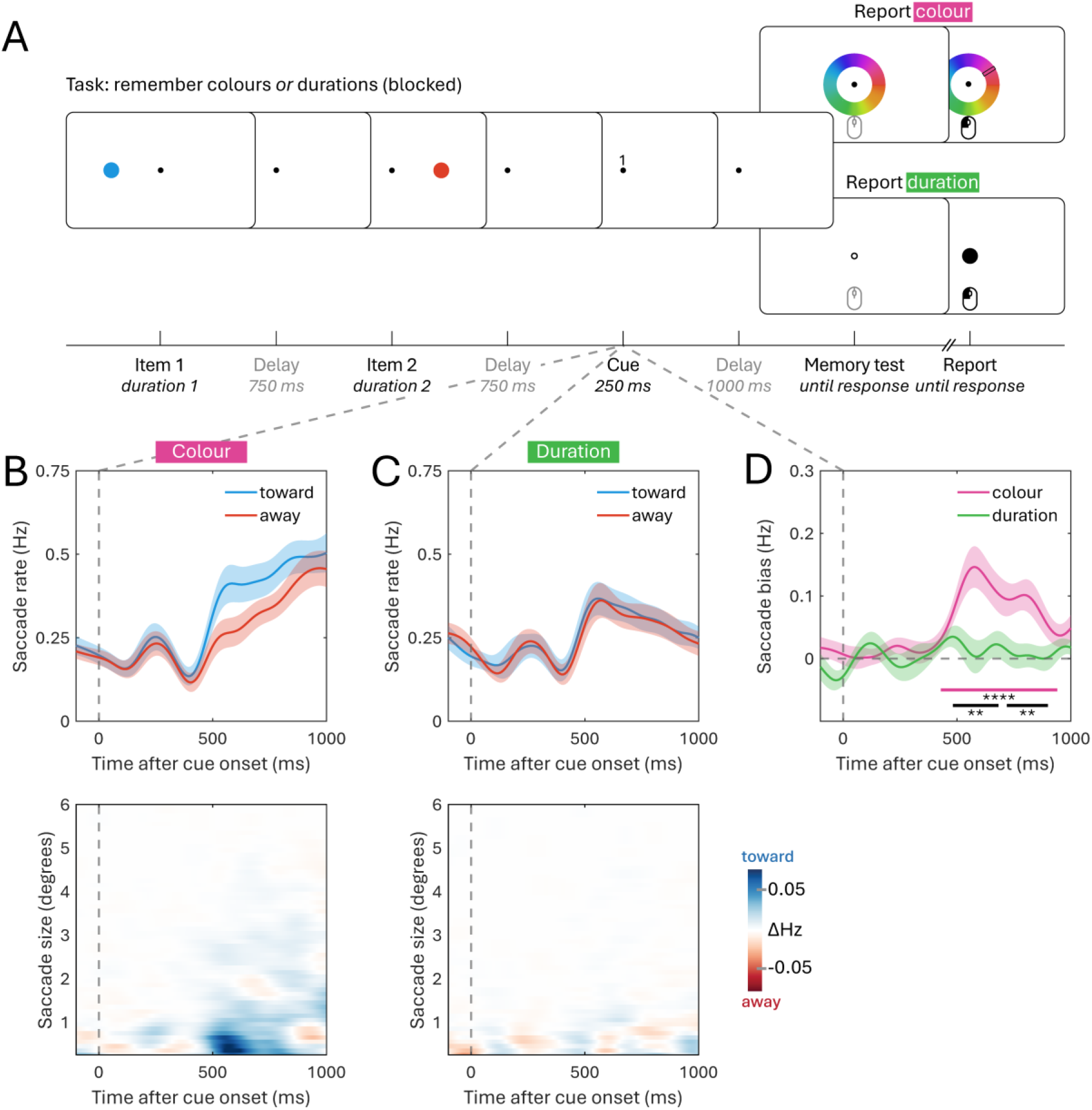
Spatial organisation is retained when memorising the colour, but not the duration of visual items in working memory. (A) Schematic of the working-memory task. Participants were presented two visual items of varying colours and durations, after which an order cue indicated which item was the target for an ensuing reproduction report. Two versions of this task were performed that used the exact same memory items and cues but varied which feature was relevant for report: in one session participants reproduced the colour of the cued item, in the other session they reproduced the duration of the cued item. In both sessions, responses were given with the mouse. Colours shown serve only as an example, for the experimental colours, see the methods section. (B) Eye-tracking data from the colour-report session. The top panel shows the time courses of saccade rates toward and away from the original location of the cued memory item; the bottom panel shows the time course of the spatial saccade bias (toward vs. away), as a function of saccade size. (C) Eye-tracking data from the duration-report session. The top panel shows the time courses of saccade rates toward and away from the original location of the cued memory item; the bottom panel shows the time course of the spatial saccade bias (toward vs. away), as a function of saccade size. (D) Overlay of time courses of the spatial saccade bias during selection from memory (toward – away), for both the colour (pink) and duration (green) conditions. All time courses indicate mean values, with shaded areas indicating the standard error of the mean. Horizontal lines indicate significant clusters (cluster-based permutation test; (Maris & Oostenveld, 2007), with the black horizontal line denoting the significant difference in the spatial saccade bias between the colour and duration sessions. Stars indicate the following levels of significance: *: p< 0.05, **: p <0.01, ***: p < 0.001, ****: p < 0.0001.

Performance in both sessions confirmed that participants relied on the cue. The average reproduction error for colour trials was 16.01±4.04 (M±SD) degrees (reports were made on a 360-degree colour wheel) and for duration trials 317.64±52.52 ms. Both observed errors were significantly lower than what would be expected if participants randomly reported the colour or duration of either of the two presented items on each trial (colour: t(23)=74.77, p<0.0001, d=20.84; duration: t(23)=59.71, p<0.0001, d=17.55; see the methods for details). This confirms that participants successfully used the cue to select the target memory item in both the colour and duration sessions.

To track the potential spatial organisation of the two items in working memory, we analysed directional biases in microsaccades during the post-cue selection period. This bias in microsaccades has recently been established as a robust peripheral index of internal attentional shifts among items held within the spatial organisation of working memory (van Ede, 2026).

We first investigated whether accessing colour, an inherently non-spatial feature of a visual item, would nevertheless lead to this previously established directional bias in microsaccades toward the cued memory item. **Figure 1B (top panel)** shows the time-resolved saccade rates following cue onset in the colour session, separated by whether saccades were directed towards or away from the cued memory item’s incidental location at encoding. We observed a clear and significant directional bias: the rate of saccades made towards the original location of the cued memory item was significantly higher than the rate of saccades made away from it (**Fig. 1D**: cluster p<0.0001). An analysis of this directional bias as a function of saccade sizes confirmed that this directional effect was principally driven by saccades in the “micro”-range (**Fig. 1B, bottom panel**). As the spatial bias was exclusively defined in memorised visual space, the items’ spatial organisation must have been retained while colour was memorised, despite colour being a purely non-spatial feature and the items’ locations never being tested.

Next, we used the exact same marker to evaluate whether this spatial organisation was also retained when tasked to remember the duration of the very same visual items, following the exact same cues. **Figure 1C (top panel)** shows the time-resolved saccade rates in the duration session. Strikingly, and in direct contrast to the colour session, accessing the cued memory item’s duration did not similarly trigger a directional microsaccade bias: there was no significant difference between the rates of saccades made towards versus away from the cued item’s original location (**Fig. 1D**: cluster p=0.219). **Figure 1C (bottom panel)** shows this difference rate split out across saccade sizes, demonstrating further that no specific saccade-size range exhibited a systematic directional bias.

Critically, a direct comparison showed that the directional microsaccade bias was significantly stronger during item selection in the colour session than in the duration session, yielding two adjacent clusters (cluster p=0.003, and cluster p=0.006; black horizontal lines in **Fig. 1D**).

Together, these results provide clear evidence for a dissociation in how the colour and duration of visual items are maintained. Specifically, colour information remains bound to the item’s spatial location at encoding, as evidenced by the spatial microsaccade bias. The absence of this microsaccade bias when retaining an item’s duration suggests that duration information is accessed in a functionally distinct way: without similarly utilising the item’s original spatial location.

## Discussion

Our results reveal a clear dissociation in the organisation of working memory for the colour versus duration of visual objects. As shown by spatial biases in microsaccades, colour information remains bound to the items’ incidental locations at encoding, but this same marker of spatial organisation was absent for duration information. This dissociation cannot be attributed to task non-compliance, stimulus confounds, or measurement insensitivity: behavioural performance confirmed cue usage across both sessions; we used the exact same stimuli, cues, and task procedure across both sessions; our microsaccade marker reliably indexed spatial organisation for colour and, instead of hinging on a null result, we report a statistically significant difference when memorising colour versus duration. Together, these findings show that the usage of space for organising non-spatial information in working memory is not universal, but feature-specific.

We tracked spatial organisation implicitly through directional biases in microsaccades when accessing the colour or duration of a cued visual memory item. Ample prior studies have demonstrated that the direction of microsaccades is biased towards spatially attended locations (e.g., Engbert & Kliegl, 2003; Hafed & Clark, 2002) and to the original location of memorised items, as was shown more recently (for a recent overview and relation to complementary findings in the literature see: van Ede, 2026). In our design, items were presented sequentially and cued by order, meaning items never competed in space during encoding. Furthermore, participants performed memory reports around central fixation. Any directional bias following the cue must therefore reflect an internally maintained spatial organisation of working memory. The clear spatial bias observed during colour recall proves that our marker is capable of indexing the spatial organisation of non-spatial features in a non-spatial task. This provides relevant context for interpreting the lack of the same bias during the session in which, instead of colour, item duration was task relevant.

We chose to directly compare the colour and duration of visual items because both are non-spatial features. Our colour results align with a large, well-established literature on other visual features, such as orientation (de Vries & van Ede, 2024; Draschkow et al., 2022; Poch et al., 2017; Schneegans & Bays, 2017), shape (Kuo et al., 2009; A. Treisman & Zhang, 2006), and real-world object identity (Liu & van Ede, 2025; Wang & Ede, 2024). In the majority of these studies, these features have been shown to rely on the spatial organisation of memory, even when item location is never asked about. In contrast, relatively few studies to date have studied working memory for visual duration, let alone the organisational principles that support it. Our findings show that duration does not abide by the same principle of spatial organisation, which highlights the importance of further investigating the mechanisms supporting working memory for visual duration.

While we reveal a clear dissociation between how duration and colour information are organised, our data do not reveal how duration information *is* organised in working memory.

Rather, our findings demonstrate that it does not share the spatial organisation characteristic of colour and other previously tested visual features. Spatial information might still be preserved alongside duration information, but in a format decoupled from the oculomotor system and therefore immeasurable through our microsaccade marker. Yet, even if duration information is retained in a spatial format not visible through the microsaccade marker, our primary conclusion holds: the underlying organisational formats for colour and duration must be qualitatively and structurally distinct. Beyond this, the task-dependent nature of our findings also add to a growing body of studies that show flexible reformatting of working-memory representation, depending on the anticipated task (e.g., Cao & Deouell, 2025; Kwak & Curtis, 2022; Liu et al., 2026; Serences et al., 2009; Yang et al., 2026).

The sensory recruitment hypothesis (Ester et al., 2009; Harrison & Tong, 2009; Phylactou et al., 2022; Roelfsema & de Lange, 2016; Serences, 2016; Sreenivasan et al., 2014; but see also: C. S. Adam et al., 2022; Xu, 2017) provides a viable explanation for the divergent spatial organization of memory for colour versus duration. In this framework, maintaining sensory information in working memory relies on the same neural regions and networks that initially process perceptual input. Consequently, the maintenance of colour information would rely more heavily on early, retinotopically mapped visual areas, where spatial position is an intrinsic organisational principle (Groen et al., 2022). Activating these areas to access a colour trace may inevitably reactivate the complementary spatial location. Duration, by contrast, is likely a higher-order property that may lack a dedicated, low-level retinotopic receptor surface.

Accordingly, the maintenance of duration information might be mediated by higher-order brain areas and networks (Centanino et al., 2026; Coull et al., 2008; Harvey et al., 2020), where access to duration traces does not recruit the early retinotopic visual maps that may be a prerequisite for observing the directional microsaccade bias during selection from working memory.

Our results corroborate the earlier findings from Kruijne et al. (2021) who provided initial and complementary evidence that duration information might be maintained in working memory without a spatial organisation. We also extend their work in at least four ways. First, we presented durations using stable visual items, instead of displaying them as the time between two briefly presented visual flashes. Second, our task likely demanded more precise memory retention by employing a continuous-report paradigm. Third, we complement the prior EEG findings during working-memory maintenance by studying a complementary microsaccade-based marker of the spatial organisation of working memory during memory selection. Finally, by directly comparing the spatial organisation of duration to that of colour, we could demonstrate a statistically significant difference between them, rather than relying exclusively on a null effect.

Even though the exact organisation of duration information in working memory remains an open question, our data reveal that its organisational architecture is fundamentally distinct from that of colour. Future research could employ more direct neural measurements to verify whether duration information is indeed maintained and accessed within more frontal, non-retinotopically organised brain regions. Moreover, with the putative lack of a spatial scaffold to organise duration information, future research should determine what alternative organisational principle keeps the duration of individual objects in working memory separate and accessible for guiding behaviour.

We conclude that working memory for object duration and colour do not share the same spatial organisation, indicating that the cognitive architecture of visual working memory is not one-size-fits-all. Although space remains a canonical organising principle, the organisation of working memory is ultimately feature-specific.

## Methods

All experimental procedures were reviewed and approved by the ethics committee of the Vrije Universiteit Amsterdam (VCWE-2022-117R1). Each participant provided written informed consent prior to participation and was reimbursed with 12.50 euros/hour or participation credits.

### Participants

All participants were recruited from the Vrije Universiteit Amsterdam. A sample size of twenty-four was set a priori, as based on previous publications from our lab with similar experimental designs and outcome measures (e.g., van Ede et al., 2019; Liu et al., 2022; Wang & Ede, 2025; de Vries & van Ede, 2025). While in these prior studies we often used a sample size of twenty-five, here we settled on twenty-four as the closest even number that enabled full counterbalancing of the order of the colour and duration sessions that were central to this experiment. After collecting the a priori determined sample size, we realised a mistake had been made in the counterbalancing, leading to thirteen participants starting with the colour session, instead of twelve. Consequently, the first participant was replaced to reach the final sample size of twenty-four participants with properly counterbalanced session orders (age range: 18-26; 23 women, 1 man; 23 right-handed).

### Task and procedure

Participants performed two near-identical versions of a working-memory task (**Fig. 1A**), that only differed in which feature of visual items was instructed to be memorised and subsequently tested at the end of a working-memory delay. The task always consisted of two visual items presented sequentially, each of which had a random colour and was presented for a variable duration. Across two consecutive sessions, participants were instructed to reproduce either the colour (colour session) or the duration (duration session) of one of these items at the end of the working-memory delay. Because the relevant response feature (colour or duration) was blocked into two separate sessions, participants knew even before the start of a trial whether the colour or duration of an item had to be remembered. Half of the participants started with the colour reproduction session, while half of the participants started with the duration reproduction session. Each session contained 10 blocks of 40 trials each. Before starting each session, participants practiced the session-specific response and task until they felt acquainted with the task.

As shown in **Figure 1A**, the first item was followed by a 750 ms delay, after which the second item was shown. This was followed by a second 750 ms delay. After this, an order cue (“1” or “2”) appeared above the central fixation dot for 250 ms, indicating which item’s colour or duration had to be reproduced at the end of the trial. The cues were 100% reliable. After cue offset, there was a final delay of 1000 ms before the participants could start their response. First (“1”) and second (“2”) cues were equally likely. The cued item was equally likely to have been presented to the left or to the right of the fixation dot, and the cued item was equally likely to have been shown for a short or a long duration. The exact duration of the cued item was randomly drawn from a short or a long range, where short durations were randomly drawn from any of the 600 possible durations between 200 to 800 ms, and long durations from any of the 600 possible durations between 1200 to 1800 ms. If the cued item was shown for a short duration, then the other item was shown for a long duration, and vice versa. The colour of both items was always randomly drawn from all 360 possible hues, with the constraint that the two items could not have the same colour.

After the final delay, participants could start their reproduction response. In colour sessions, the report was prompted by the presentation of the colour response wheel (**Fig 1A, top right**).

The colour response wheel was always shown with a random orientation offset for each trial, so participants could not prepare the required action in advance. Once participants moved the mouse, a custom cursor appeared on the colour wheel which participants could move around for as long as they wanted, until they clicked the left mouse button, which locked in the currently selected colour as the response. In duration sessions, the report was prompted by changing the colour of the fixation dot to black (**Fig 1A, bottom right**). Participants reproduced the memorised duration of the cued item by clicking the left mouse button, which made a colourless version of the original item appear centrally. Participants kept the left mouse button pressed down for the duration they intended to reproduce. The accompanying item stayed on screen until the left mouse button was released. In both session types, feedback was shown after the response. Colour feedback was given as a score between 0 to 100, where 100 meant there was a 0° distance between the target colour and the responded colour, and 0 meant the maximum distance of 180°. Duration feedback was given as the signed difference between the responded duration and the target duration, rounded to milliseconds.

### Experimental set-up

Both tasks were programmed in Python 3.10.18 using the PsychoPy library (46) to generate the stimuli. Participants were seated approximately 70 cm in front of a 24-inch LCD monitor, with a 1920 × 1080 pixel resolution and a 239 Hz refresh rate.

The presented items were always circles with a diameter of 2 degrees visual angle, presented at 6.5 degrees visual angle from central fixation (from the centre of the screen to the centre of either item; the distance to the edge of either item was always 5.5 degrees visual angle). For the colour of the items, the HSV colour system was used. All 360 possible hues (H) were used while the saturation (S) and value (V) were constant at, respectively, 20% and 50%. The fixation dot was a near-white (rgb: 234, 234, 234) circle with a diameter of 0.2 degrees visual angle, but changed to black (rgb: 0, 0, 0) in duration sessions to indicate the response could start. The cue (rgb: 255, 255, 255) was presented 0.3 degrees visual angle above central fixation. The colour wheel shown at response in colour session was a doughnut shape with an outside diameter of 12 degrees visual angle and an inside diameter of 9 degrees visual angle. During the duration response, the original visual item (2 degrees visual angle in diameter) was shown centrally, but always in white (rgb: 255, 255, 255), so that the original colour was never reshown. This ensured participants were required to use the order cue to know which item’s duration to reproduce.

### Eye-tracking acquisition and preprocessing

Horizontal and vertical gaze positions of the participant’s right eye were continuously sampled throughout the experiment using an EyeLink 1000 Plus at a rate of 1000 Hz. The eye-tracker was positioned approximately 10 cm in front of the monitor, and 60 cm away from the eyes.

Participants used a chinrest to minimise head movements. Prior to recording, the eye tracker was calibrated and validated using the built-in HV9 calibration module. The eye tracker was always re-calibrated in between the two sessions (after 10 blocks), and could additionally be re-calibrated in each break between consecutive blocks, if the signal was deemed of too poor quality.

Following acquisition, eye-tracker datafiles were converted from their original Eyelink Data Format (.edf) to an ASCII text file (.asc), and analysed in MATLAB R2024b using a combination of the Fieldtrip analysis toolbox (Oostenveld et al., 2011) and custom code. Custom code was used to detect blinks and remove them from the signal for 100 ms before and after each blink (by replacing these data with NaNs). After blink removal, data were epoched relative to cue onset.

### Saccade detection and visualisation

Saccades were detected using a previously established and validated velocity-based method (Liu et al., 2022). In this approach, saccades are detected by finding the samples where the 2-dimensional gaze velocity exceeds a trial-based threshold. Gaze velocity was defined as the derivative of gaze position, after smoothing in the temporal dimension with a Gaussian-weighted moving mean filter with a 7-ms sliding window (using the built-in MATLAB function ‘smoothdata’). The velocity threshold was set to 5 times the median velocity in any given trial, which is consistent with earlier work (de Vries & van Ede, 2025; Liu & van Ede, 2025; van Harmelen & van Ede, 2025; Wang & Ede, 2025). A minimum delay of 100 ms between successive saccades was imposed to avoid counting the same saccade multiple times.

Saccade size (in degrees) and saccade direction (left/right) were calculated by estimating the difference between the pre-saccade gaze position (−50 to 0 ms before threshold crossing) and the post-saccade gaze position (50 to 100 ms after threshold crossing). Depending on the horizontal (left/right) direction of the saccade and the original left/right location of the cued memory item, saccades were classified as ‘toward’ or ‘away’. After detecting and classifying the saccade directions, the time courses of saccade rates (in Hz) were obtained using a sliding time window of 100 ms, advancing in steps of 1 ms.

To also investigate the size of the saccades that contributed to our findings, without imposing an inherently arbitrary threshold, we additionally decomposed saccade rates into a time-size representation, showing the time courses of saccade rates as a function of the saccade size (as in van Ede, 2026). For this, we used successive saccade-size bins of 0.5 degrees visual angle in steps of 0.1 degree.

### Statistical analysis

Prior to analysing the performance data, we z-scored the decision times for each participant and removed all trials with an absolute z-score greater than 3 from the performance analysis. Decision times were defined as the time between response prompt and response initiation.

To statistically evaluate the performance data to verify the use of the cue, we compared the observed reproduction errors to the reproduction errors that would have occurred if the order cue had not been used. We followed the same procedure for colour and reproduction reports. Expected errors for the scenario in which the cues were ignored were calculated by assuming perfect memory reports, that were centred on the correct (cued) item in half the trials and on the incorrect (uncued) item in the other half of the trials. Accordingly, the expected error in this scenario would be the distance between the two items in a given trial, divided by two (as in half the trials the error would be zero, and in the other half it would be the full distance between the two items). We averaged this value across all trials, to obtain the hypothetical best possible performance in the scenario where the cue was ignored. For the statistical comparison between the observed and these hypothetical reproduction errors, paired Student’s t-tests were employed. Effect sizes were quantified using Cohen’s d.

To statistically evaluate the time series saccade data, we employed a non-parametric cluster-based permutation approach (Maris & Oostenveld, 2007). Using this approach we statistically evaluated the time courses of the spatial saccade bias by comparing the rate of toward versus away saccades. We did this separately for the colour and duration conditions. We used the same statistical approach to also directly compare the time courses of the observed spatial saccade biases (quantified as the difference between toward and away saccades) between the colour and duration conditions. All three permutation analyses were conducted on the time period from 0 to 1000 ms after cue onset, using Fieldtrip with default clustering settings (grouping adjacent time points that were significant in a mass-univariate comparison and summing their t-values to arrive at a cluster size). Permutation distributions of the largest cluster size were acquired by randomly permuting the condition labels of each participant’s trial-averaged time course data (i.e. randomly flipping the labels of toward/away saccades or of the colour/duration conditions) 10,000 times and identifying the size of the largest clusters observed in these randomised data after each permutation. The p-values of the clusters observed in the original data were calculated as the proportion of random permutations where the largest cluster was equal to or larger than the cluster that we observed in the original (unpermuted) data.

## Acknowledgements

This research was supported by an NWO Vidi Grant by the Dutch Research Council (grant number 14721), and an ERC Starting Grant from the European Research Council (MEMTICIPATION, 850636) to F.v.E. In addition, we thank Laurie Rol for collecting the data for this experiment, and we thank Christos Dalamarinis and Emma van den Brink for their contributions to the earlier projects that eventually led to the project we present here.

## Data, materials and software availability

All raw data and analysis code will be made public prior to publication. The code for the experimental task is already publicly available on [GitHub](https://github.com/annavanharmelen/Microsaccade-bias-colour-vs.-duration-experiment).

